# EV-Tracer enables lineage-resolved detection and molecular profiling of extracellular vesicle-associated signals in cancer–fibroblast co-culture

**DOI:** 10.64898/2026.08.25.746941

**Authors:** Yutaka Naito, Chiaki Hori, Keisuke Yoshida, Takanori Amano, Masakazu Yashiro, Kazuyoshi Yanagihara, Kazufumi Honda

**Affiliations:** Department of Molecular Prevention, Institute for Advanced Medical Sciences, Nippon Medical School, 1-1-5, Sendagi, Bunkyo-ku, Tokyo, 113-8602, Japan; Department of Oral Pathobiological Science and Surgery, Tokyo Dental College, 2-9-18, Kandami-saki-Cho, Chiyoda-Ku, Tokyo 101-0061, Japan; Next Generation Human Disease Model Research Team, BioResource Research Center, RIKEN, 3-1-1 Koyadai, Tsukuba, Ibaraki, 305-0074, Japan; Department of Molecular Oncology and Therapeutics, Graduate School of Medicine, Osaka Metro-politan University, 1-4-3 Asahimachi, Abeno-ku, Osaka 545-8585, Japan; Division of Rare Cancer Research, National Cancer Center Research Institute, 5-1-1 Tsukiji, Chuo-ku, Tokyo, 104-0045, Japan; Department of Molecular Prevention, Graduate School of Medicine, Nippon Medical School, 1-1-5, Sendagi, Bunkyo-ku, Tokyo, 113-8602, Japan

**Keywords:** Extracellular vesicles, EV-Tracer, CD63, Tumor microenvironment

## Abstract

Extracellular vesicles (EVs) facilitate intercellular communication by transferring diverse bioactive molecules from donor to recipient cells. However, EVs released by distinct cellular lineages become difficult to distinguish when mixed in multicellular experimental models, limiting the analysis of how cell–cell interactions affect EV-associated molecular profiles. To address this, EV-Tracer, a CD63-based dual-fluorescence tracing and capture system for detecting, isolating, and profiling lineage-associated EV fractions, is developed. Achilles or mScarlet is inserted into the small extracellular loop of CD63, enabling tracer-specific EV detection by digital counting, antibody-based isolation, and live-cell visualization. Exploratory EV RNA sequencing suggests that physical cell–cell contact is associated with distinct EV RNA profiles, including interferon-related signals, which are supported by targeted cellular and EV-associated RNA analyses. EV-Tracer provides a practical framework for investigating lineage-associated EV dynamics and molecular signals in mixed-cell systems.

## Introduction

Tumors comprise not only cancer cells but also non-malignant stromal cell types, such as immune cells, endothelial cells, smooth muscle cells, pericytes, mesothelial cells, mesenchymal stem cells (MSCs), and resident fibroblasts. Intercellular communication between cancer cells and these stromal cells is pivotal for cell proliferation, cancer cell invasion, therapeutic resistance, and metastasis (1–3). Soluble factors, direct cell–cell or cell–extracellular matrix contact, and extracellular vesicles (EVs) mediate these interactions. Through these interactions, tumor and stromal cells dynamically change their phenotypes and functions over time.

EVs are lipid bilayer membrane vesicles that contain various bioactive molecules, such as mRNAs, microRNAs (miRNAs), lipids, metabolites, and proteins. These molecules in EVs can be transferred to recipient cells in proximal tissues and distal organs, where they affect diverse cellular phenotypes (3–5). However, conventional EV studies typically rely on monoculture or bulk EV analysis. Monoculture-based EV analyses retain information about the producing cell type but cannot reproduce interaction-dependent changes in cellular phenotypes. Conversely, bulk EV analysis from mixed-cell cultures can detect EV changes associated with cell-cell interactions but loses lineage information. Although fluorescent and biochemical EV-labeling methods have enabled visualization of EV processes (6, 7) and flow cytometry separation of EVs, few approaches enable EV labeling, selective isolation, quantitative detection, and lineage-resolved molecular profiling in mixed-cell models. This limitation makes it difficult to assess how different interaction states, such as indirect, non-contact co-culture and direct, contact co-culture, affect EV-associated signals from specific cell lineages.

To address this limitation, we developed EV-Tracer, a dual-color, CD63-based tracing and capture system. We introduced fluorescent proteins into the small extracellular loop of CD63 to display distinct tracer epitopes on CD63-positive vesicles. We generated the 44As3 gastric cancer cell line expressing CD63exAchilles (CD63exAch) and the iNF-58 stomach fibroblast line expressing CD63exmScarl (CD63exScarl). We evaluated EV-Tracer using ExoCounter digital EV counting, antibody-conjugated magnetic bead isolation, and live-cell imaging. We then applied EV-Tracer to mono-, indirect-, and direct-co-culture conditions and performed exploratory RNA profiling of the recovered lineage-associated EV fractions. This approach enabled us to examine whether EV-associated RNA signals differed across cellular lineages and according to the physical interaction state of the co-culture.

## Methods

### Cell cultures

The 44As3 cell line and the iNF-58 immortalized human normal stomach fibroblast line were used, as described previously (8). The HCT116 cell line was purchased from ATCC (CCL-247). Briefly, all cell lines were cultured in high-glucose Dulbecco’s modified Eagle medium containing sodium pyruvate (DMEM; Thermo Fisher Scientific, Waltham, MA, USA), 10% FBS and 1% penicillin-streptomycin (Thermo Fisher Scientific). The A549 cell line, used as the source of CD63 cDNA, was obtained from the Japanese Collection of Research Bioresources (JCRB) Cell Bank and cultured under the same conditions. For direct co-culture experiments, the 44As3 cell line (2 × 10^5^ cells) was seeded into the same well as iNF-58 (2 × 10^5^ cells). For indirect co-culture experiments, the 44As3 cell line (2 × 10^5^cells) was seeded into the top of the Transwell insert. The iNF-58 fibroblast line (2 × 10^5^ cells) was cultured in the lower compartment of a 6-well plate, separated by a 0.4-μm pore size filter (Corning Life Science, Tewksbury, MA, USA), allowing humoral factors to pass through. For the monoculture condition, 2 × 10^5^ cells were seeded into each well. Monoculture, indirect co-culture, and direct co-culture experiments were performed in high-glucose DMEM containing sodium pyruvate (Thermo Fisher Scientific), 1% FBS, and 1% penicillin-streptomycin (Thermo Fisher Scientific). For the monoculture reference condition, cell-conditioned medium (CCM) collected separately from 44As3 and iNF-58 in monocultures was combined in equal volumes before analysis. The total amount of CCM under each culture condition was adjusted to match the others, then applied to the subsequent experiments.

### Plasmids

The AAVS1 hPGK-PuroR-pA donor was a gift from Rudolf Jaenisch (Addgene plasmid # 22072; RRID: Addgene_22072) (9) and was used as the backbone vector for both CD63exAch and CD63exScarl, which were constructed as EV-Tracer plasmids. The Achilles/pRSETB was provided by the RIKEN BRC through the National BioResource Project of the MEXT, Japan (cat. RDB15982) (10). The mScarlet vector was purchased from Addgene (Plasmid #141153). The human CD63 coding sequence was amplified from cDNA prepared from total RNA isolated from the A549 cell line. The complete CD63 coding sequence was verified by Sanger sequencing and was confirmed to be identical to the reference human CD63 sequence (NM_001780.6). Achilles was inserted into the first small extracellular loop after codon 43, as previously reported (11). mScarlet was also inserted at the corresponding position with the RDPPV linker. The cloning primers are listed in Table S1. For the CRISPR/Cas9 system, pX330-U6-Chimeric_BB-Cbh-hSpCas9 was a gift from Feng Zhang (Addgene plasmid #42230; RRID: Addgene_42230) (12). The hPGK-PuroR-pA cassette was inserted into the PmeI site of this plasmid, and then a guide RNA targeting the AAVS1 locus was subsequently introduced (13). The resulting plasmid was designated pX330-puro-AAVS1 and used in this study.

### Transfection and stable line construction

Cell transfection was performed using Lipofectamine 2000 Reagent (Thermo Fisher Scientific) according to the manufacturer’s instructions. Briefly, cells were seeded to 50-70% confluence the day before transfection. Then, 400 ng each of EV-Tracer plasmids and pX330-puro-AAVS1 were mixed with 100 μL of Opti-MEM (Thermo Fisher Scientific) and Lipofectamine reagent, then combined with 500 μL of growth medium. After two days of incubation, the medium was replaced with growth medium containing 2-8 μg/mL puromycin, and the transfected cells were selected for seven days.

### Extracellular vesicle (EV) isolation

The cells were washed with PBS (−), and the culture medium was replaced with Advanced DMEM medium (Thermo Fisher Scientific) for the HCT116 cell line. After 48 h of incubation, the CCMs were collected and centrifuged at 2,000 × g for 10 min at 4 °C. To thoroughly remove cellular debris, the supernatant was filtered through a 0.22-μm filter (Merck Millipore, Billerica, MA, USA). CCMs of HCT116 cell lines were then used for EV isolation. The EV purification was performed as described previously (8). Briefly, CCMs of HCT116 cell lines expressing either EV-Tracer or AAVS1 control were ultracentrifuged at 100,000 × g for 70 min at 4 °C using a SW41Ti rotor (Optima XE-90, Beckman Coulter, Brea, CA, USA). The pellets were washed with PBS (-), ultracentrifuged again under the same conditions, and resuspended in PBS (−). The isolated EVs were visualized using a transmission electron microscope (TEM) outsourced to Tokai Electron Microscopy, Inc. (Nagoya, Japan). The samples were adsorbed onto formvar film-coated copper grids and stained with 2% phosphate-tungstic acid solution (pH 7.0) for 30 s. The grids were observed by a transmission electron microscope (JEM-1400Plus; JEOL Ltd., Tokyo, Japan) at an acceleration voltage of 100 kV. The image of EVs was acquired with a CCD camera (EM-14830RUBY2; JEOL Ltd.). The protein concentration of the putative EV fraction was determined by the Micro BCA Protein Assay Kit (Thermo Fisher Scientific). To determine the size distribution of the EVs, nanoparticle tracking analysis was performed using the NanoSight LM10 system (NanoSight Ltd., Amesbury, UK) on samples diluted 100-fold with PBS (-).

To prepare proteins for Western blot, CCMs were centrifuged at 2,000 × g for 10 min at 4 °C, and the supernatants were filtered through a 0.22-μm filter. Tracer-associated EV fractions were isolated using a modified antibody-affinity capture protocol based on the Biotin Capture Magnetic Beads and Immobilising/Washing Buffer supplied in the MagCapture Exosome Isolation Kit PS Ver. 2 (FUJIFILM Wako Pure Chemical Corporation, Osaka, Japan). Briefly, 60 μL of the streptavidin-magnetic beads were incubated with biotinylated anti-Achilles (anti-GFP antibody, clone FM264G, Cat: 338002, Lot. No. B365717, Biolegend, San Diego, CA, USA) or anti-mScarlet (anti-RFP antibody, clone 8E5.G7, Cat: 200-301-379, Lot. No. 47969, Thermo Fisher Scientific) or IgG antibody (clone RTK2758, Cat: 400501, Lot. No. B403407, Biolegend) for 10 min at room temperature in the Immobilising/Washing buffer, with the beads inverted. Antibody biotinylation was performed using the Biotin Labeling Kit - NH2 (Dojindo, Kumamoto, Japan) according to the manufacturer’s instructions. After three washes with Immobilising/Washing buffer, antibody-conjugated magnetic beads were incubated with 500 μL of CCM samples for 24 h at 4 °C. After three washes with Immobilising/Washing buffer, an appropriate lysis buffer was added to isolate total RNA or protein. To prepare EV samples for RNA-seq and qRT-PCR, CCMs containing 1% FBS were centrifuged at 2,000 × g for 10 min at 4 °C. The supernatants were filtered through a 0.22-μm filter and ultracentrifuged at 100,000 × g for 70 min at 4 °C using a SW41Ti rotor. The pellets were washed with PBS (−), ultracentrifuged again under the same conditions, and resuspended in PBS (−). The isolated EVs were subjected to antibody-conjugated magnetic bead isolation as described above.

### Quantification of EV counts by ExoCounter

The CCM samples and EV fractions from HCT116, 44As3, and iNF-58 cell lines with EV-Tracer were analyzed using the ExoCounter system (JVCKENWOOD Corp., Kanagawa, Japan) (14) according to the manufacturer’s instructions. Briefly, an optical disc with a removable well plate containing 16 wells for sample injection (CB398470, JVCKENWOOD Corp.) was coated with 5 μg/mL anti-Achilles (anti-GFP antibody, clone FM264G, Cat: 338002, Lot. No. B365717, Biolegend) or anti-mScarlet antibody (anti-RFP antibody, clone 8E5.G7, Cat: 200-301-379, Lot. No. 47969, Thermo Fisher Scientific) in PBS for 1 h at 37 °C. Anti-CD9 or anti-CD63 preconjugated discs were purchased from JVCKENWOOD Corp. (AB188730 and BW875160). After washing with PBS containing 0.05% Tween 20 (PBS-T), the discs were incubated with a blocking solution (1% BSA in PBS-T). The CCM samples or EV fractions (50 μL) were incubated for 2 h at 37 °C with shaking. After washing three times with PBS-T, diluted anti-CD9-or anti-CD63-antibody-conjugated beads (100× dilution in PBS-T) were incubated for 90 min at 37 °C with shaking. Each well was first washed five times with PBS-T, then three times with deionized water. The disc was dried at 37 °C for 15 min, and then Achilles- or mScarlet-positive and CD9-positive EVs were quantified with the ExoCounter. The corresponding background count was subtracted from the sample count in each experiment. Negative background-subtracted values were retained for statistical analysis and interpreted as background-level signals. For CCMs, DMEM containing 10% FBS was used as a negative background for EV-Tracer characterization (Fig. 2B, 2C, 3C, 3D, and S1B), whereas DMEM containing 1% FBS was used as a negative background for comparisons among monoculture, indirect co-culture, and direct co-culture conditions (Fig. 4D). PBS-T was used as a negative background for isolated EVs.

### Immunoblot analysis

For immunoblot analysis, whole-cell lysates and EV lysates from magnetic beads were prepared using Mammalian Protein Extract Reagent (M-PER; Thermo Fisher Scientific) with a protease inhibitor cocktail (SIGMA, P8340). The whole-cell lysates and isolated EVs were solubilized in Laemmli sample buffer by boiling, then loaded onto NuPAGE Bis-Tris Mini Protein Gels (4-12%, Thermo Fisher) and subjected to electrophoresis (150 V). The proteins were transferred to a polyvinylidene difluoride membrane (Millipore). After blocking in 10% BSA/PBS (Albumin, from Bovine Serum, Fraction V pH 7.0, FUJIFILM Wako Pure Chemical Corporation), the membranes were incubated for 1 h at room temperature with primary antibodies, which included anti-CD63 (Clone 8A12, Cat: SHI-EXO-M02, Lot. No. 21K24CB, dilution 1: 200, Cosmo Bio, Tokyo, Japan), anti-CD9 (Clone 12A12, Cat: SHI-EXO-M01, Lot. No. 26G24CB, dilution 1: 200, Cosmo Bio), anti-actin (Clone AC-15, Cat: AB6276, Lot. No. 1107472-2, dilution 1: 5,000, Abcam), and anti-GM130 (Clone EP892Y, Cat: AB52649, Lot. No. 1151072-2, dilution 1: 1000, Abcam). Secondary antibodies (horseradish peroxidase-linked anti-mouse IgG, Cat: NA9310, Lot. No. 18236611 or horseradish peroxidase-linked anti-rabbit IgG, Cat: NA9340, Lot. No. 18130507, Cytiva, Tokyo, Japan) were used at a dilution of 1: 5,000. The membrane was then exposed to SuperSignal West Dura Extended Duration Substrate (Thermo Fisher Scientific).

### Cell proliferation assay

HCT116 cell lines with EV-Tracers and AAVS1 control (2000 cells) were seeded into each well of a 96-well plate. Cell viability was determined on the indicated days using the CellTiter-Glo 2.0 Cell Viability Assay (Promega, Madison, WI, USA) according to the manufacturer’s instructions, and luminescence was measured using a SynergyTM H4 Microplate Reader (BioTek, Winooski, VT, USA). Measurements were performed using separate matched wells at Day 0 and Day 2. Technical replicate wells (6 to 12 wells) were averaged at each time point, and an ATP-based growth index was calculated as the mean Day 2 luminescence divided by the mean Day 0 luminescence. Three independent experiments were performed.

### Microscopic images in vitro

The fluorescence and bright-field images of CD63exAch and CD63exScarl cells were acquired using the BZ-X810 microscope (KEYENCE, Osaka, Japan). For time-lapse imaging, 44As3 with CD63exAch and iNF-58 with CD63exScarl were seeded into the same well of the Ibidi μ-Plate 24-well plate at 2 × 10^4^ cells each and incubated for 24 h. The cells were then scanned at 60x objective magnification, and time-lapse images were recorded every 5 min over 145 min (30 time points) using the time-lapse imaging unit of the BZ-X810 microscope.

### RNA extraction and quantitative RT-PCR (qRT-PCR)

Total RNA was extracted from cultured cells using the miRNeasy Plus Mini Kit (Qiagen, Hilden, Germany) according to the manufacturer’s protocols. For total RNA extraction from EV fractions, the miRNeasy Plus Micro Kit (Qiagen) was used. For mRNA expression by qRT-PCR analysis, complementary DNA (cDNA) was generated from total RNA using SuperScript IV VILO Master Mix (Thermo Fisher Scientific). For EV-associated RNA analysis, QuantAccuracy RT-RamDA cDNA Synthesis Kit (TOYOBO, Osaka, Japan) was used. Real-time PCR was then performed in triplicate with cDNA using Fast SYBR Green Master Mix (Thermo Fisher Scientific). The data were collected and analyzed using the QuantStudio 3 Real-Time PCR System (Thermo Fisher Scientific). All mRNA quantification data from cultured cells were normalized to the expression of β-actin (ACTB). All primer sequences are listed in Table S1. For EV-associated RNA analysis, equal volumes of starting CCMs were processed using identical ultracentrifugation, affinity capture, and RNA extraction procedures. RNA was eluted in the same final volume for all samples, and equal aliquots of the resulting RNA eluates were used for reverse transcription. IFITM3 RNA abundance was normalized to ACTB and calculated using the 2^−ΔΔCt^ method. For cellular qRT-PCR analysis, the 44As3 and the iNF-58 cells were cultured separately in monoculture, harvested separately after 48 h, and pooled as cell populations before RNA extraction. For the indirect co-culture condition, 44As3 cells from the upper insert and iNF-58 cells from the lower compartment were harvested separately after 48 h and pooled before RNA extraction. For the direct co-culture condition, the mixed-cell population was harvested together. Total cellular RNA was then extracted from the combined 44As3 and iNF-58 cell populations under each condition. Equal amounts of total RNA were used for reverse transcription in all samples.

### RNA-seq and bioinformatics

Transcriptomic library construction and bulk RNA-seq analysis of total RNA from magnetic bead-isolated EV samples in monoculture and in indirect and direct co-cultures (n=1 per reporter−condition combination) were performed by Rhelixa Corporation (Tokyo, Japan). RNA quality and concentration were determined using a Bioanalyzer. Equal amounts of input RNA were used for cDNA library preparation with the SMART-Seq mRNA HT LP kit (Takara Bio Inc., Shiga, Japan) according to the manufacturer’s instructions. The sequencing was conducted on an Illumina NovaSeq X Plus under the following conditions: 150 bp paired-end (PE150), 6 Gb per sample, and 40 million reads (20 million read pairs) per sample.

FASTQ files were pseudo-aligned, and transcript abundances were estimated using kallisto (version 0.46.2) against the Ensembl human GRCh38 cDNA reference. Transcript-level abundance estimates were imported into R (version 4.4.3) using the tximport package. Gene-level estimated counts were analyzed using edgeR. Trimmed mean of M-value (TMM) normalization was applied, and log2 counts per million were calculated using the cpm function with a prior count of 1. These normalized expression values were used for subsequent exploratory analyses.

Principal component analysis was performed on the 1,000 most variable genes with a mean log2-CPM greater than 1 across the six libraries using the prcomp function in R. For pathway analysis, the Hallmark gene set collection was obtained from the Molecular Signatures Database using the msigdbr package. Gene sets represented by fewer than 10 genes in the EV RNA-sequencing expression matrix were excluded. Gene set variation analysis was performed using the GSVA package with a Gaussian kernel. For heatmap visualization, GSVA scores were standardized to row-wise z-scores across the displayed samples and truncated to the range of −1.2 to 1.2.

### Statistical analysis

Values are presented as means ± s.d. for biological replicates. Statistical analyses between the two groups were performed using Welch’s t-test. For multiple comparisons within each experiment, one-way ANOVA with Tukey’s HSD test was used. A p-value less than 0.05 was considered significant. All statistical analyses were performed using R software version 4.4.3.

### Artificial intelligence-assisted tools

ChatGPT (OpenAI, San Francisco, CA, USA) assisted in generating and refining part of the R code for RNA-seq data analysis. The authors critically reviewed, modified as needed, and executed all AI-assisted code. The authors made all analytical decisions, interpreted the results, and drew the scientific conclusions. During manuscript preparation, the authors used DeepL (DeepL SE, Cologne, Germany) and ChatGPT (OpenAI) to improve the manuscript’s grammar, spelling, and readability. A professional English-language editor subsequently proofread the manuscript.

## Results

### Development of a dual-color, CD63-based EV-Tracer system

To distinguish EV populations associated with different cellular lineages, two CD63-based fluorescent tracers were designed (Fig. 1A). CD63 was chosen as the membrane scaffold for EV-Tracer because it is a widely used sEV-associated tetraspanin, is preferentially enriched in intraluminal vesicles of late endosomal multivesicular bodies (MVBs), and has been extensively used to label and track MVB trafficking and CD63-positive sEV secretion (11, 15). A fluorescent protein can be inserted into the first small extracellular loop of CD63, where it becomes exposed on the outer surface of EVs released from MVBs (11). Based on this topology, we inserted Achilles or mScarlet, bright, monomeric fluorescent proteins, into the small extracellular loop of CD63 to generate CD63exAch and CD63exScarl, respectively. The fluorescent domains, located on the extracellular surface of CD63-positive vesicles, are intended to enable fluorescence-based tracking in live cells and to facilitate antibody-mediated detection and isolation of labeled EVs using specific antibodies against Achilles or mScarlet fluorescent proteins.

**Fig. 1.**
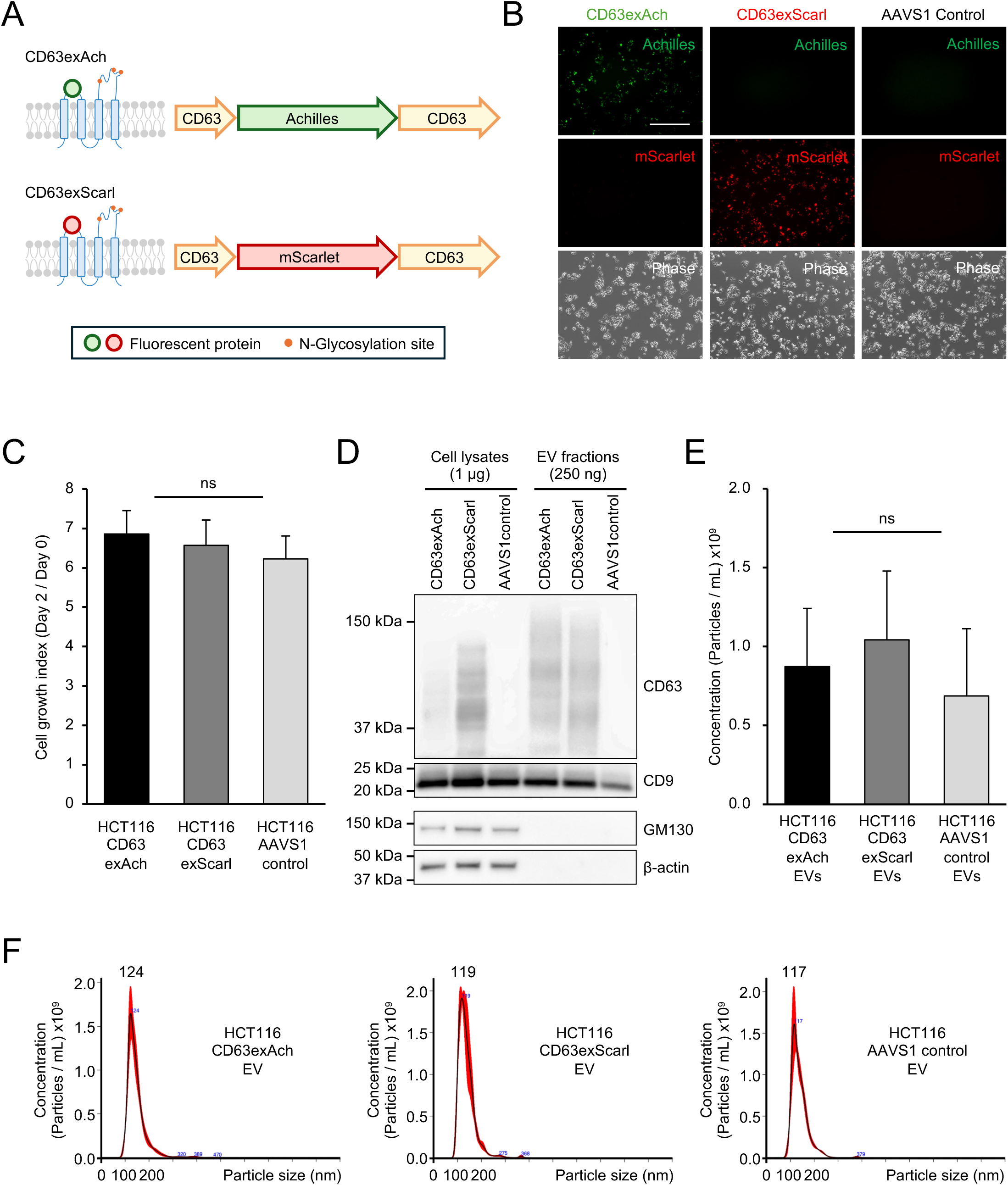
Effects of CD63exAch and CD63exScarl on cellular behavior and the characteristics of secreted EVs. (A) Schematic illustration of the EV-Tracer, CD63exAch and CD63exScarl. Achilles or mScarlet was inserted into the small extracellular loop of CD63, thereby displaying the fluorescent reporter on the extracellular surface of CD63-positive vesicular structures. Predicted N-glycosylation sites are indicated as orange dots. (B) Representative Achilles, mScarlet, and phase-contrast images of HCT116 cell lines stably expressing CD63exAch, CD63exScarl, or the AAVS1 control construct. Scale bar, 400 μm. (C) ATP-based growth indices of HCT116 CD63exAch, HCT116 CD63exScarl, and HCT116 AAVS1 control cells. The growth index was calculated as the mean Day 2 luminescence divided by the mean Day 0 luminescence. Data are shown as means ± s.d. from three independent experiments. The indicated comparisons were performed using Welch’s t-test. ns, not significant. (D) Representative immunoblot analysis of CD63, CD9, GM130, and β-actin in whole-cell lysates and isolated EV fractions from the indicated HCT116 cell lines. (E) Particle concentrations of ultracentrifuged EV preparations determined by nanoparticle tracking analysis (NTA). Data are shown as means ± s.d. from three independent EV preparations. The indicated comparisons were performed using Welch’s t-test. ns, not significant. (F) Representative particle-size distributions of EV preparations from HCT116 CD63exAch, CD63exScarl, and AAVS1 control cells. Modal particle diameters are indicated above the respective distributions.

We selected HCT116 colorectal cancer cells for initial characterization of EV-Tracer because this cell line had previously been used to evaluate EV detection (14). We generated stable HCT116 cell lines by integrating the CD63exAch construct, the CD63exScarl construct, or the control vector into the AAVS1 safe-harbor locus. The fluorescence of Achilles and mScarlet was observed in the corresponding vector-expressing cells (Fig. 1B). Transmission electron microscopy showed vesicular structures with comparable morphology in EV preparations from CD63exAch and AAVS1 control cells (Fig. S1A). Expression of both CD63exAch and CD63exScarl did not grossly alter cell proliferation or cellular morphology in the HCT116 cell line (Fig. 1B and 1C). Immunoblotting detected CD63 and CD9 in the isolated EV fractions from all three HCT116 cell lines (Fig. 1D and Fig. S3). In contrast, GM130 and β-actin were detected only in cell lysates (Fig. 1D). Nanoparticle tracking analysis did not identify a substantial difference in particle concentration or size distribution (Fig. 1E and 1F). Consistent with these findings, ExoCounter, a digital EV counting method (14), also showed no significant differences in EV counts using the CD63–CD9 and CD9–CD9 antibody combinations between CCMs from HCT116 CD63exAch and HCT116 AAVS1 control cells. In contrast, counts using the CD63–CD63 antibody combination were higher in CCM from HCT116 CD63exAch cells (Fig. S1B). These data suggested that EV-Tracer expression might increase CD63 abundance on individual EV particles or the proportion of CD63-positive EVs, but might not affect the total number of EVs secreted by cells.

Next, to investigate the specificity of EV-Tracer-based detection, CCM from the engineered cell lines was incubated with anti-Achilles or anti-mScarlet antibody-conjugated discs of the ExoCounter system, each combined with anti-CD9 antibody-conjugated beads (Fig. 2A). Anti-Achilles or anti-mScarlet antibodies, combined with anti-CD9 antibody-conjugated beads, selectively detected EV particles in the corresponding EV-Tracer-positive CCMs (Fig. 2B and 2C) and isolated EV fractions (Fig. 2D and 2E). Although we detected a small number of nanoparticles in CCMs from the AAVS1 control and non-corresponding tracer-expressing cells, signals from these cell lines remained at background levels. These results demonstrated tracer-dependent detection with minimal detectable cross-reactivity between the two antibody assays.

**Fig. 2.**
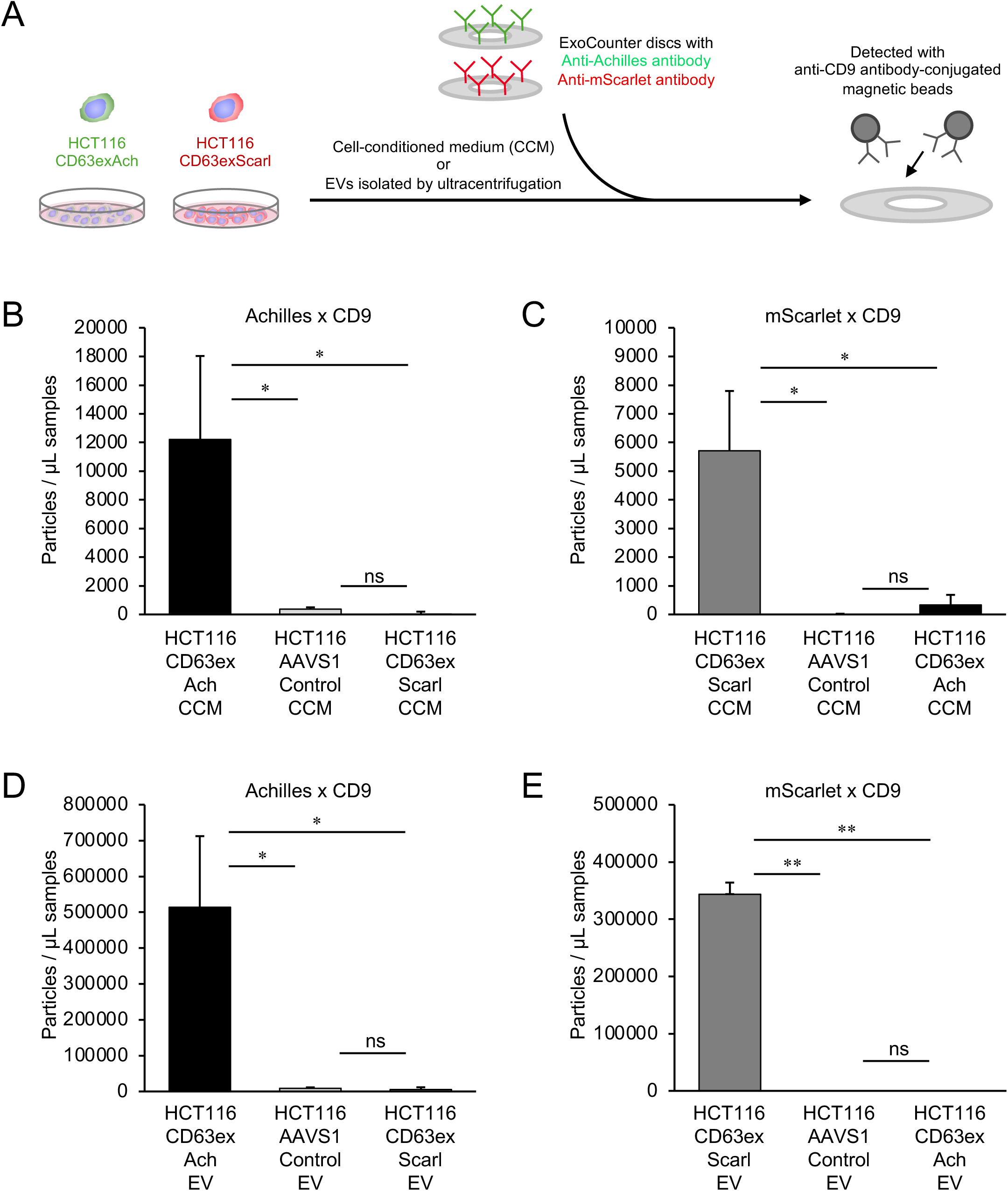
Tracer-specific detection of CD63exAch and CD63exScarl-expressing EV particles by ExoCounter. (A) Schematic representation of the ExoCounter assay for investigating the number of CD63exAch- and CD63exScarl-expressing EV particles. Cell-conditioned medium (CCM) or isolated EV fractions from HCT116 CD63exAch, CD63exScarl, or AAVS1 control cells were applied to optical discs coated with anti-Achilles or anti-mScarlet antibody and detected using anti-CD9 antibody-conjugated magnetic beads. (B, C) Achilles–CD9 (b) and mScarlet–CD9 (c) ExoCounter measurements in CCMs from the indicated HCT116 cell lines. Matched 10% FBS medium background counts were subtracted. (D, E) Achilles–CD9 (d) and mScarlet–CD9 (e) measurements in the corresponding isolated EV fractions. PBS-T background counts were subtracted. Data are shown as means ± s.d. from three independent experiments. Statistical comparisons were performed using Welch’s t-test. *p < 0.05; **p < 0.01; ns, not significant.

We next interrogated whether these EV-Tracers could be used for cell-mixture experiments. CD63exAch was introduced into the 44As3 gastric cancer cell line (16), and CD63exScarl was introduced into the iNF-58 stomach fibroblast line (8) (Fig. 3A). Fluorescence microscopy showed intracellular distributions of Achilles and mScarlet signals in the respective cell lines, consistent with localization to CD63-positive endosomal and vesicular compartments (Fig. 3B). As in HCT116 cell lines, the anti-Achilles and anti-mScarlet assays detected EV particles in the corresponding tracer-positive CCMs (Fig. 3C and 3D).

**Fig. 3.**
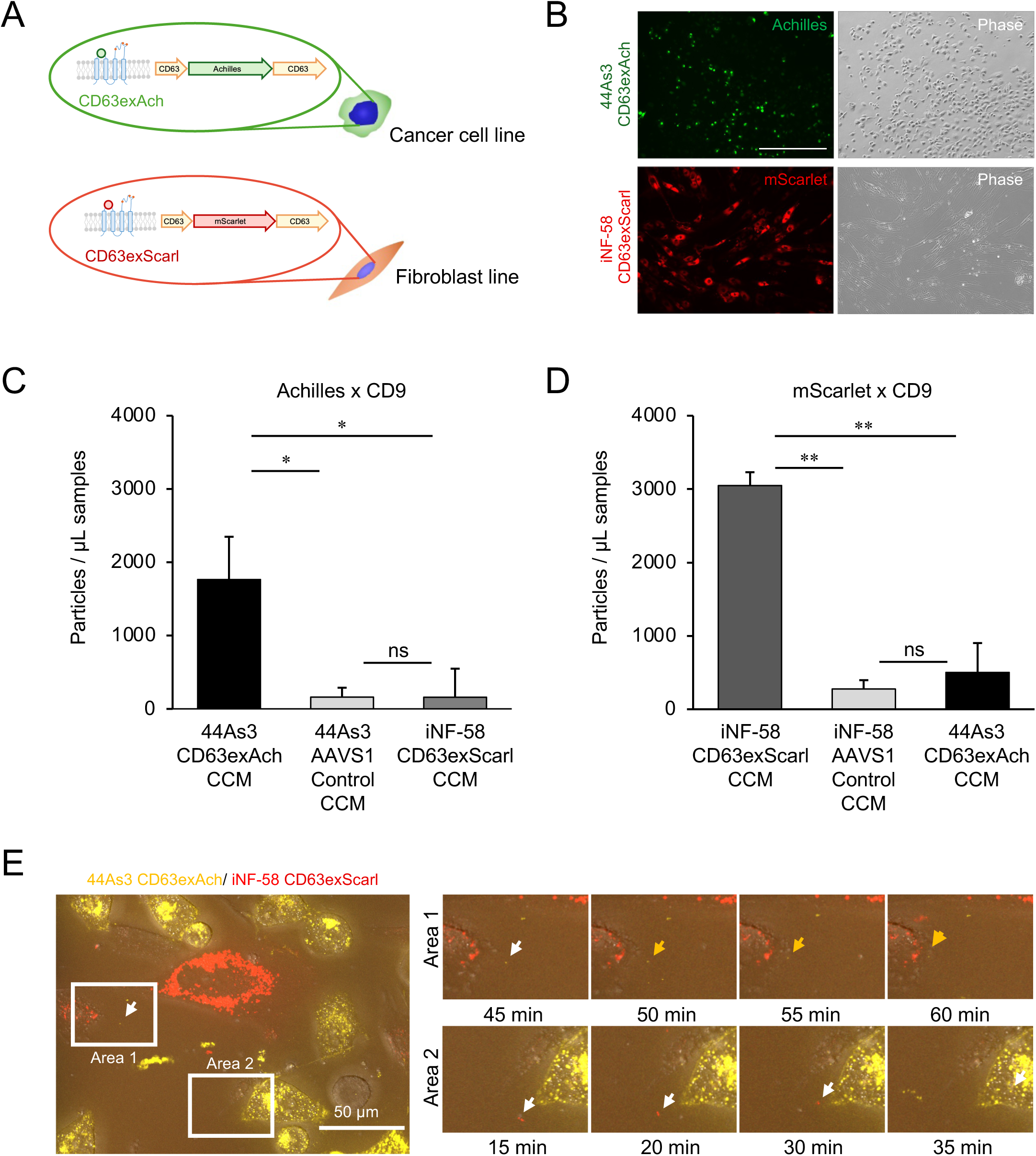
Establishment of lineage-assigned EV-Tracer expressing cell lines and live-cell visualization of reporter-positive puncta. (A) Schematic representation of the tracer assignment used in the heterotypic co-culture model. CD63exAch is expressed in the 44As3 gastric cancer cell line, whereas CD63exScarl is expressed in the iNF-58 stomach fibroblast line. (B) Representative fluorescence and phase-contrast images of 44As3 CD63exAch and iNF-58 CD63exScarl cells. Scale bars, 500 μm. (C, D) Achilles–CD9 (c) and mScarlet–CD9 (d) ExoCounter measurements in cell-conditioned medium (CCM) from the indicated tracer-expressing and AAVS1 control cell lines. Matched 10% FBS medium background counts were subtracted. Data are shown as means ± s.d. from three independent experiments. Statistical comparisons were performed using Welch’s t-test. *p < 0.05; **p < 0.01; ns, not significant. (E) Left: Representative time-lapse images of directly co-cultured 44As3 CD63exAch and iNF-58 CD63exScarl cells. Right: A series of higher-magnification images of Area 1 and Area 2 in the left image. Arrows indicate representative Achilles- or mScarlet-positive puncta that changed position during imaging. Times indicate elapsed time from the beginning of the displayed sequence. Scale bar, 50 μm. The complete video image is provided as Video S1.

We also investigated the utility of these two individual EV-Tracers microscopically. Live-cell fluorescence imaging of co-cultured 44As3-CD63exAch cells and iNF-58-CD63exScarl fibroblasts showed dynamic movement of Achilles- and mScarlet-positive puncta between neighboring cells (Fig. 3E and Video S1). These observations support the use of EV-Tracer to visualize tracer-positive vesicular structures in mixed-cell environments.

### EV-Tracer enables isolation and quantification of lineage-associated EV fractions in co-culture

We next tested whether EV-Tracer could recover lineage-associated EV fractions from a shared CCM (Fig. 4A). 44As3-CD63exAch cancer cells and iNF-58-CD63exScarl fibroblasts were cultured together under direct co-culture conditions. We then incubated CCMs with magnetic beads conjugated to anti-Achilles or anti-mScarlet antibodies. The recovered fractions were analyzed by immunoblotting for the EV-associated markers CD63 and CD9 (Fig. 4B and Fig. S4). CD63 and CD9 were detected with both anti-Achilles and anti-mScarlet antibody-conjugated beads, but not with control IgG-conjugated beads (Fig. 4B and Fig. S4). Together with the antibody-specificity results from monoculture CCM (Fig. 3C and 3D), these findings showed that EV fractions associated with each fluorescent tracer could be isolated from a common co-culture medium. We then used EV-Tracer to compare EV-associated signals across three culture conditions: monoculture, indirect co-culture, and direct co-culture (Fig. 4C). In the indirect co-culture condition, cancer cells and fibroblasts were separated by a permeable insert, allowing exchange of soluble factors without direct physical contact. In the direct co-culture condition, two cell types were cultured in the same wells to allow direct physical contact. After two days, we analyzed CCMs using ExoCounter assays with CD63–CD9, Achilles–CD9, and mScarlet–CD9 antibody combinations. Total CD63–CD9 signals and lineage-associated tracer signals showed only modest differences among mono-, indirect-, and direct-co-culture conditions (Fig. 4D). Thus, direct physical contact might not increase EV-associated signals during the two-day culture period, at least in CD63- and CD9-double-positive EVs derived from 44As3 and iNF-58. EV uptake and subsequent recycling or re-release cannot be excluded in mixed-cell systems. Thus, the recovered populations are lineage-associated EV fractions, rather than definitive cell-of-origin EV populations.

**Fig. 4.**
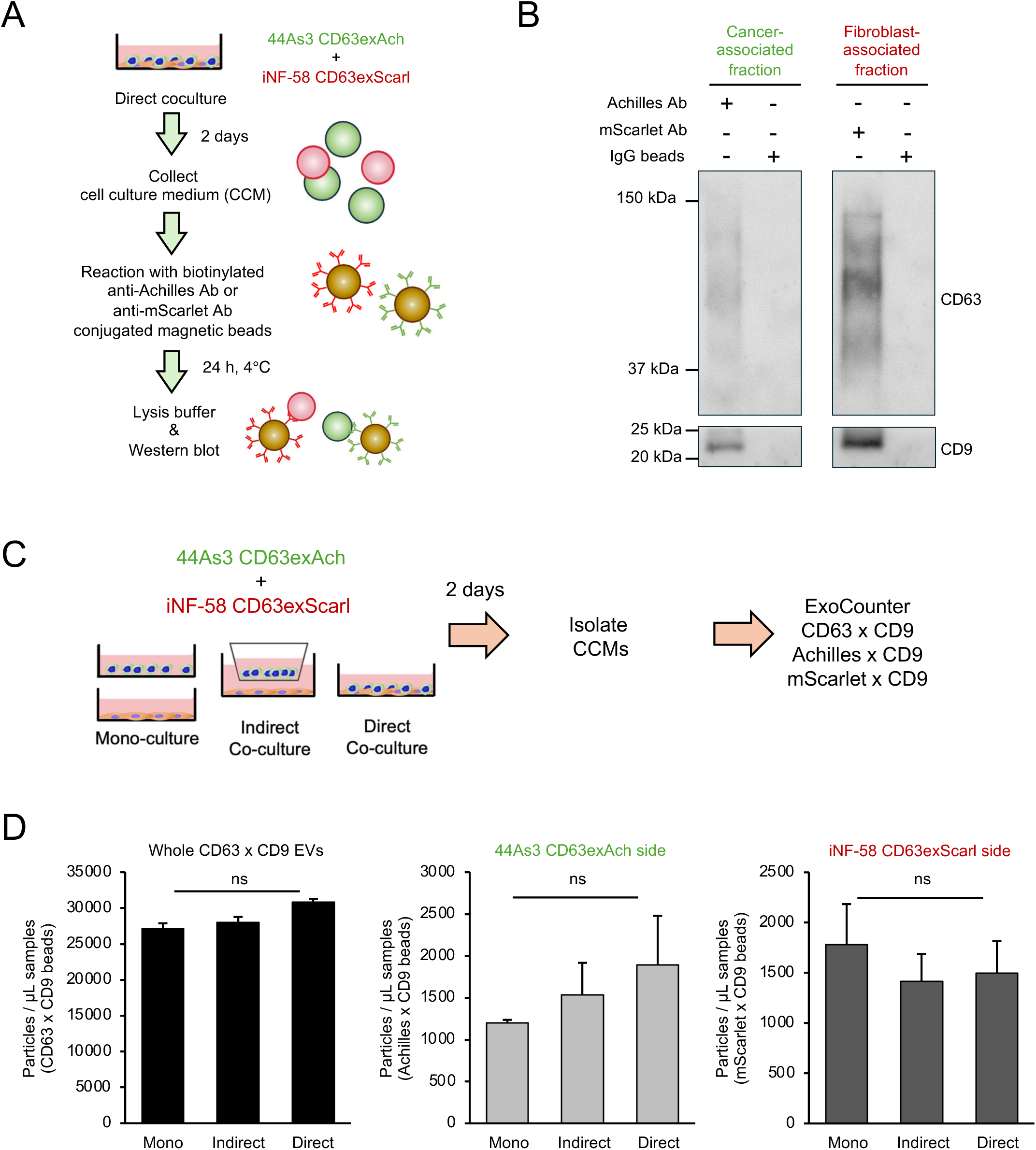
Dual-tracer affinity capture and quantitative detection of lineage-associated EV fractions from cancer–fibroblast co-cultures. (A) Experimental workflow for reporter-specific magnetic-bead capture. Cell-conditioned medium (CCM) from the direct-co-culture condition of 44As3 CD63exAch and iNF-58 CD63exScarl cells was incubated with anti-Achilles-or anti-mScarlet-conjugated magnetic beads for 24 h at 4 °C, followed by lysis and immunoblot analysis. (B) Immunoblot analysis of CD63 and CD9 in the recovered fractions from direct co-culture CCM using magnetic beads conjugated to anti-Achilles, anti-mScarlet, or control IgG antibodies. (C) Schematic representation of ExoCounter analysis in the monoculture, indirect-co-culture, and direct-co-culture conditions. For the monoculture reference, equal volumes of CCM collected separately from 44As3 CD63exAch and iNF-58 CD63exScarl monocultures were combined before analysis. CCMs were collected after two days and analyzed using CD63–CD9, Achilles–CD9, and mScarlet–CD9 ExoCounter assays. (D) Total CD63–CD9-positive particle counts and Achilles–CD9- and mScarlet–CD9-positive reporter-associated particle counts under the three culture conditions. Matched 1% FBS medium background counts were subtracted. Data are shown as means ± s.d. from three independent biological experiments. The indicated comparisons were performed using Welch’s t-test. ns, not significant.

### Exploratory RNA profiling identifies contact-associated signals in lineage-associated EV fractions

To test whether EV-Tracer could support downstream molecular profiling of the isolated EV fractions, we maintained 44As3-CD63exAch cells and iNF-58-CD63exScarl fibroblasts under monoculture, indirect co-culture, or direct co-culture conditions for two days. Because the amount of EV-associated RNA recovered directly from CCMs was limited, we first isolated EV fractions by ultracentrifugation. Then we separated them using anti-Achilles- or anti-mScarlet-conjugated magnetic beads. RNA recovered from the isolated fractions was then sequenced (Fig. 5A). Principal component analysis showed separation of the Achilles-associated and mScarlet-associated EV RNA profiles and further suggested variation corresponding to the monoculture, indirect co-culture, and direct co-culture groups (Fig. 5B). These findings can support the feasibility of recovering molecularly distinguishable lineage-associated EV fractions.

**Fig. 5.**
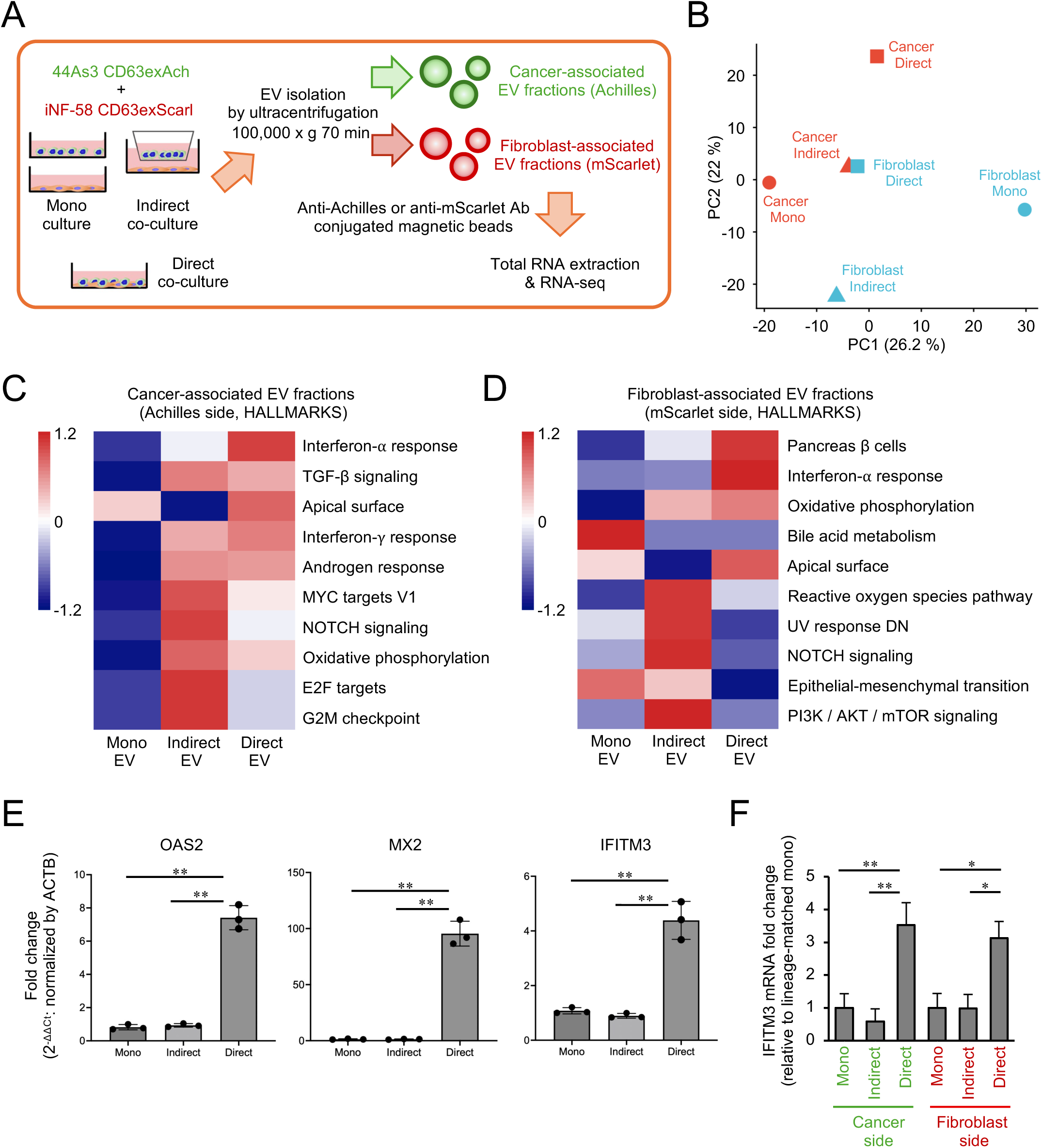
Exploratory RNA profiling of lineage-associated EV fractions under different cancer– fibroblast interaction states. (A) Experimental workflow for RNA profiling of lineage-associated EV fractions. 44As3 CD63exAch and iNF-58 CD63exScarl cells were maintained under monoculture, indirect co-culture, or direct co-culture conditions. EV-containing fractions were isolated by ultracentrifugation and subsequently separated using anti-Achilles- or anti-mScarlet-conjugated magnetic beads before RNA extraction and sequencing. (B) Principal component analysis of the six EV RNA-sequencing libraries using the 1,000 most variable genes with a mean log2-CPM greater than 1. The percentages of variance explained by PC1 and PC2 are indicated. (C, D) The five Hallmark gene sets with the highest GSVA scores in the direct-co-culture sample and the five with the highest scores in the indirect-co-culture sample were selected and displayed together. When the same gene set appeared in both lists, the next-highest-ranked non-duplicated gene set was selected to retain a total of 10 pathways. All GSVA results are shown in Fig. S2A. Colors indicate row-wise z-scores of GSVA scores. (E) qRT-PCR analysis of OAS2, MX2, and IFITM3 in total cellular RNA collected from the complete cultures under monoculture, indirect co-culture, and direct co-culture conditions. Expression levels of each gene were normalized to ACTB and are presented relative to monoculture. Data are shown as means ± s.d. from three independent biological experiments; individual points represent biological replicates. Statistical comparisons were performed using one-way ANOVA with Tukey’s HSD test. *p < 0.05; **p < 0.01. (F) qRT-PCR analysis of IFITM3 RNA in Achilles-associated and mScarlet-associated EV fractions. Equal starting volumes of conditioned medium were subjected to identical ultracentrifugation, affinity capture, and RNA extraction procedures. IFITM3 was normalized to ACTB and is presented relative to the corresponding lineage-matched monoculture sample, set to 1. Data are shown as means ± s.d. from three independent biological experiments. Statistical comparisons were performed using Welch’s t-test. *p < 0.05; **p < 0.01.

We therefore performed gene set variation analysis (GSVA) to visualize pathways for each condition. The relative representation of several Hallmark gene sets showed distinct descriptive patterns among the three interaction states in both cancer-associated and fibroblast-associated EV fractions (Fig. 5C and 5D). Whole Hallmark pathway profiles also differed among the three culture conditions (Fig. S2A). Notably, interferon-related gene sets showed higher scores in both EV fractions in the direct co-culture than in the corresponding monoculture or indirect co-culture fractions (Fig. 5C and 5D). cGAS-sensing and STING-IRF3 signaling via direct cell–cell interaction trigger interferon-stimulated gene (ISG) expression in both cancer cells and surrounding fibroblasts (17). We next examined representative ISGs by qRT-PCR using total cellular RNA from the combined cancer-cell and fibroblast populations. ISG expression, including OAS2, MX2, and IFITM3, was higher under direct co-culture conditions than under monoculture or indirect co-culture conditions (Fig. 5E and Fig. S2C). Consistent with these cellular findings, EV RNA-seq profiles also showed higher RNA abundance of ISGs, including IFITM3, under direct co-culture conditions (Fig. S2B). Thus, we used ISG transcripts as targets for qRT-PCR to validate our exploratory RNA-seq data of EV fractions. Of these candidates, qRT-PCR detected IFITM3 in the recovered lineage-associated EV fractions. IFITM3 mRNA abundance in EV-associated fractions was increased under direct co-culture conditions compared with monoculture and indirect co-culture (Fig. 5F). These results were consistent with the exploratory EV RNA-sequencing profiles and supported a contact-associated increase in IFITM3 mRNA abundance in the recovered EV fractions. Together, these findings demonstrate that EV-Tracer enables tracer-specific detection, isolation, and downstream molecular interrogation of lineage-associated EV fractions in mixed-cell cultures.

## DISCUSSION

In this study, we developed EV-Tracer, a dual-color, CD63-based tracing and capture system to trace information packaged into EVs under mixed-cell culture conditions. The EV-Tracer system was designed as an overexpression model in which the vectors were integrated into a safe-harbor locus. However, our characterization suggested that EV-Tracer expression had little effect on cell morphology, growth, or overall EV secretion. Furthermore, antibody-based bead isolation enables comprehensive analysis of lineage-associated EV signals and pathways under co-culture conditions. The Achilles-associated and mScarlet-associated EV fractions exhibited distinct pathway profiles, including higher interferon-response scores under direct co-culture. Independent qRT-PCR further confirmed increased IFITM3 RNA abundance in the corresponding EV-associated fractions. Total counts of CD63/CD9 double-positive EVs did not differ significantly among culture conditions after two days, suggesting that contact-associated changes in EV RNA profiles can occur without an increase in the number of these EVs. EV-Tracer exhibited intense fluorescence, enabling its use for living-cell imaging. These observations indicated that the EV-Tracer approach enabled us to examine whether EV-associated RNA signals differed across cellular lineages and according to the physical interaction state of the co-culture.

Numerous reports have highlighted the essential role of EVs in maintaining physiological homeostasis and in the progression of various diseases, including cancer. These findings have also promoted the development of novel treatment strategies and diagnostic methods for cancer (5, 18–21). Generally, EVs isolated from the CCM of monocultured cells or from body fluids have been used for functional analyses. Monoculture-derived EVs are appropriate for studying EV functions associated with a defined cell type, since they avoid contamination by EVs released from other cellular lineages. In contrast, multiple cellular lineages constitute a complex microenvironment within tumor tissues and change their phenotypes over time. Thus, the information packaged into EVs may also fluctuate during cell–cell interactions within the tumor microenvironment (22). Indeed, direct comparisons of contact and non-contact co-culture models have shown that physical cell–cell interaction can affect EV production, molecular cargo, or EV-mediated transfer (23–25). In addition, existing approaches based on engineered tetraspanins have primarily emphasized live imaging, in vivo cell-type-specific labeling, or cell-selective proteomic analysis (11, 26). For example, fluorescent CD63 reporter systems, including ExoBow (27), have enabled imaging of EV secretion and uptake in vitro (11) as well as spatial and cell-type-restricted visualization of CD63-positive exosomes in vivo (26). The Snorkel-tag, combined with StEVAC, efficiently retrieves large amounts of intact tagged EVs in vivo and in vitro (28). As a complementary approach that does not involve tetraspanin engineering, SIEVE uses cell-selective metabolic proteome labeling and bioorthogonal ligation to recover intact EVs for cell-of-origin-resolved proteomics, including from complex co-cultures (29). However, none of these approaches has enabled direct comparison of lineage-associated EV fractions from both interacting cell populations within the same heterotypic co-culture. Thus, EV-Tracer complements these approaches by integrating dual-lineage fluorescent labeling, antibody-based digital detection, dual-tracer affinity capture, and downstream RNA analysis within a shared heterotypic co-culture system.

Although EV-Tracer may help us understand new mechanisms underlying intercellular communications via EVs, the approach has several limitations. First, EV-Tracer cannot distinguish EVs directly secreted by reporter-expressing cells from reporter-positive EVs that recipient cells have internalized, repackaged, or subsequently re-released. Uptake followed by intact EV re-release or transcytosis has been demonstrated (30–32), although its prevalence and quantitative contribution in heterotypic cell co-culture conditions remain unclear. Importantly, intact EV re-release from the endosomal compartment of recipient cells may preserve the original lineage reporter and at least part of the molecular cargo, thereby retaining lineage-associated information. In contrast, membrane remodeling or repackaging within recipient cells (32) could complicate the lineage origin of EVs. Future studies, such as quantification of Achilles–mScarlet double-positive EVs, are needed to determine the extent of EV re-release and reporter mixing in the current co-culture model. Second, EV-Tracer analyses were restricted to reporter-positive, CD63-associated EV subsets and therefore did not encompass CD63-negative EV populations. In addition, the absolute abundance of CD63exAch- and CD63exScarl-associated EVs cannot be directly compared using the current antibody-based assays, because reporter expression, antibody affinity, and capture efficiency may differ between the two systems. High-sensitivity nanoflow cytometry (33) and fluorescence-activated nanoscale vesicle sorting (34–37) offer complementary advantages, enabling single-particle analysis and the potential resolution of Achilles-positive, mScarlet-positive, and double-positive EV populations. Such approaches are particularly useful for assessing reporter mixing and secondary EV processing in co-culture. However, fluorescence-based EV sorting remains constrained by instrument-dependent detection thresholds, low particle recovery, prolonged sorting, and dilution of recovered particles, which may reduce the molecular complexity available for downstream RNA analysis (36, 37). In the present study, the reporter-specific affinity capture using the EV-Tracer approach recovered lineage-associated EV fractions in a concentrated form suitable for exploratory RNA profiling. Combining single-particle nano-flow analysis with affinity-based bulk recovery may provide a more comprehensive strategy for evaluating both EV population structure and molecular cargo in future studies.

In summary, the present results demonstrate that EV-Tracer enables tracer-specific detection, isolation, and downstream molecular interrogation of lineage-associated EV fractions in mixed-cell cultures. The exploratory RNA analyses further suggest that physical cancer–fibroblast contact is reflected in EV-associated RNA signals, providing a proof-of-concept application of the platform for studying extracellular vesicle communication across defined cell-interaction states. This EV-tracing approach provides a useful platform for dissecting lineage-associated EV communication, particularly in the tumor microenvironment, and it may contribute to a better understanding of biological processes mediated by EVs.

## Additional files

Additional file 1: Supporting_information_Fig.S1_to_S4.pdf

Additional file 2: Table S1.xlsx

Additional file 3: Video S1.mp4

## Declarations

### Ethics approval and consent to participate

Not applicable.

### Consent for publication

Not applicable.

### Availability of data and materials

The raw and processed RNA-sequencing data have been deposited in the NCBI Gene Expression Omnibus under accession number GSE342194. Other data supporting our findings in this manuscript are available from the corresponding author on reasonable request.

### Authors’ contributions

Y.N. and K.H. were involved in study design, data interpretation, and writing the manuscript. Y.N. and C.H. were involved in data analysis. Y.N., Ke.Y., and T.A. established the plasmid vectors. Y.N. and C.H. were involved in sample collection. M.Y. and Ka.Y. provided research resources and supported experimental preparation. All authors critically reviewed the report, commented on drafts of the manuscript, and approved the final report.

## Acknowledgments

The authors appreciate the contribution of all members of the Department of Molecular and Cellular Medicine, Tokyo Medical University, and of the Division of the Laboratory of Integrative Oncology, National Cancer Center Research Institute, and of the Division of Interdisciplinary Genetics and Nanomedicine, Research Center for Drug Discovery, Keio University Faculty of Pharmacy, for support with EV isolation and quantification, and of the Department of Biochemistry, Kochi University Medical School for technical assistance with ExoCounter. The authors also thank Keyence Corp. for technical support with image acquisition using EV-Tracer, Meiwafosis Co., Ltd. for support with EV particle measurement, and JVCKENWOOD Corp. and Sysmex Corp. for technical support with ExoCounter. The authors also thank Forte Co., Ltd. for English proofreading. The authors are also grateful to all members of the Department of Molecular Prevention at Nippon Medical School for their stimulating discussions and technical and administrative support.

## Table of Contents

EV-Tracer enabled tracer-specific detection, affinity capture, live-cell imaging, and RNA profiling of lineage-associated extracellular vesicle (EV) fractions in mixed-cell culture conditions. In a proof-of-concept application, exploratory RNA sequencing and independent qRT PCR indicated that direct cell contact was associated with interferon-related EV RNA signals.

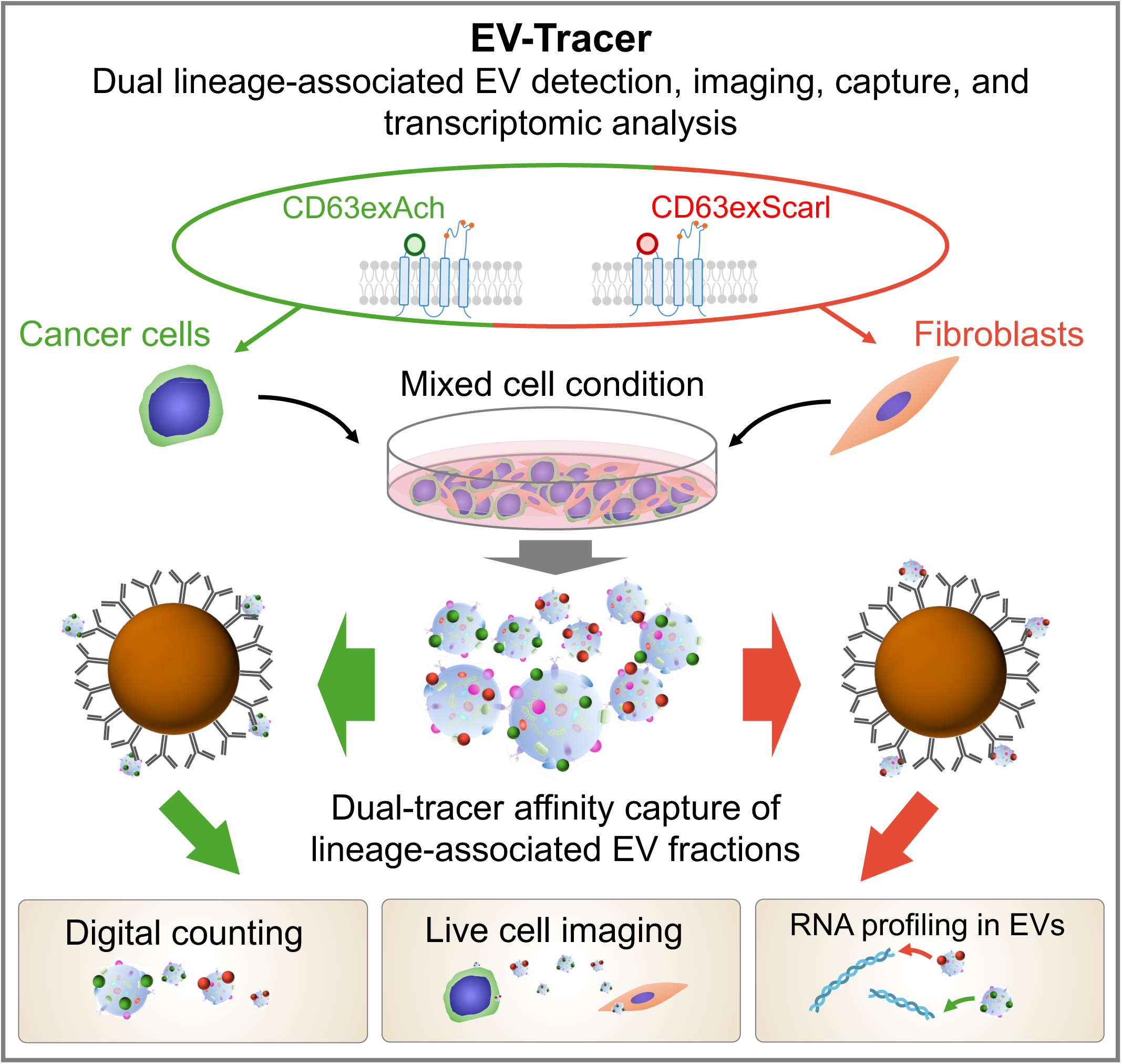

