## Supplementary information for "EV-Tracer enables lineage-resolved detection and molecular profiling of extracellular vesicle-associated signals in cancer–fibroblast co-culture"

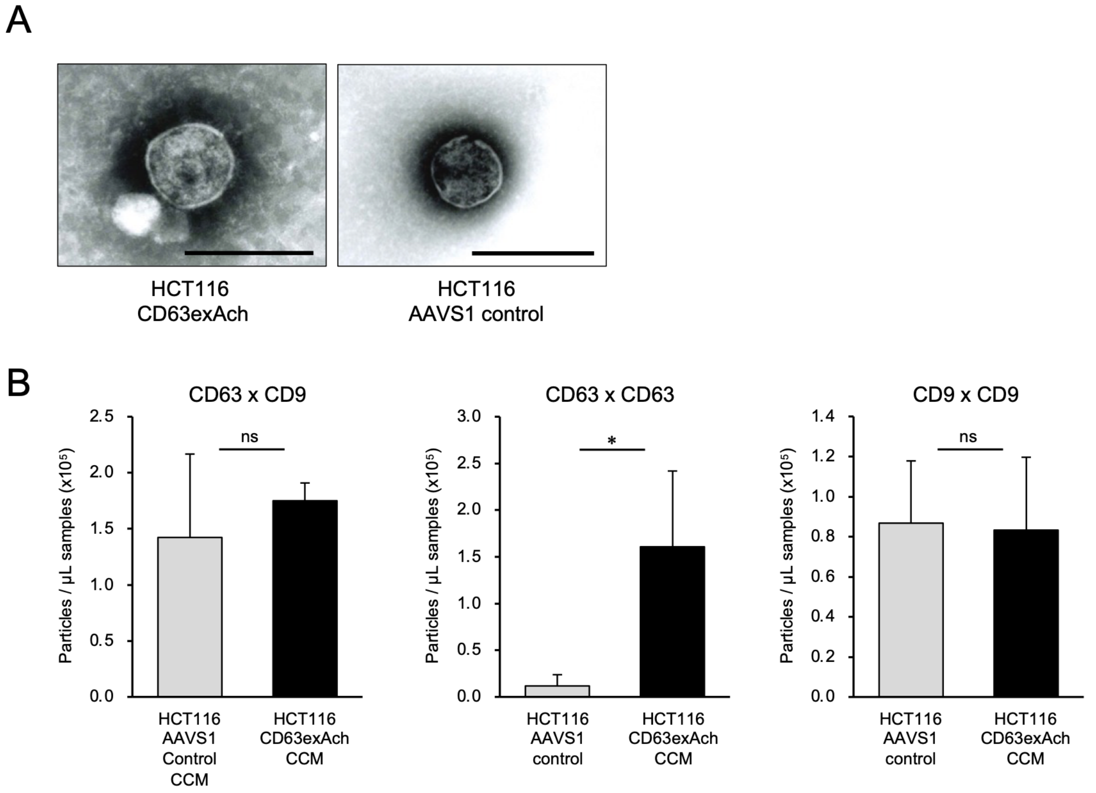


**Fig. S1 Morphological and ExoCounter characterization of EV preparations from HCT116 CD63exAch cells.**

(A) Representative transmission electron micrographs of ultracentrifuged EV preparations from HCT116 CD63exAch and AAVS1 control cells. Scale bars, 200 nm.

(B) ExoCounter measurements of CD63–CD9-, CD63–CD63-, and CD9–CD9-positive particles in cell-conditioned medium (CCM) from HCT116 CD63exAch and AAVS1 control cells. Run-specific matched medium-only background counts were subtracted. Data are shown as means ± s.d. from three independent experiments. Statistical comparisons were performed using Welch's t-test. *$p < 0.05$; ns, not significant.

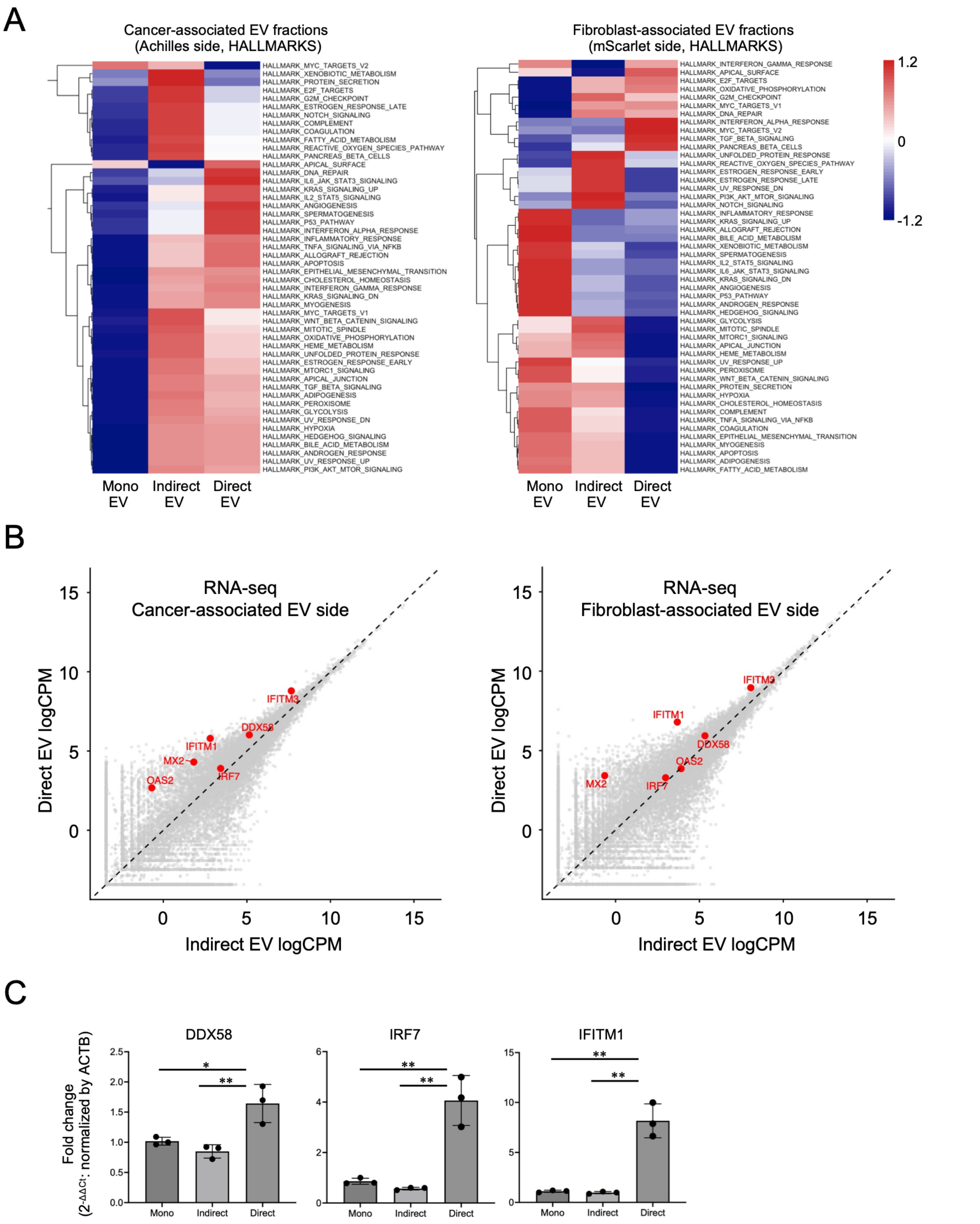


**Fig. S2 Complete Hallmark profiles and interferon-stimulated RNA patterns in lineage-associated EV fractions and corresponding cultures.**

(A) The GSVA scores for all Hallmark gene sets of each lineage-associated EV fraction in the monoculture, indirect-, and direct-co-culture conditions. Colors indicate row-wise z-scores of GSVA scores.

(B) TMM-normalized log2-CPM values from EV samples in the direct- and indirect-co-culture samples are plotted separately for the Achilles-associated (defined as cancer-associated) and mScarlet-associated (defined as fibroblast-associated) fractions.

(C) qRT-PCR analysis of DDX58, IRF7, and IFITM1 in total cellular RNA collected from the complete cultures under monoculture, indirect co-culture, and direct co-culture conditions. Expression levels of each gene were normalized to ACTB and are presented relative to monoculture. Data are shown as means ± s.d. from three independent biological experiments; individual points represent biological replicates. Statistical comparisons were performed using one-way ANOVA with Tukey's HSD test. *p $< 0.05$; **p $< 0.01$.

Fig. S3

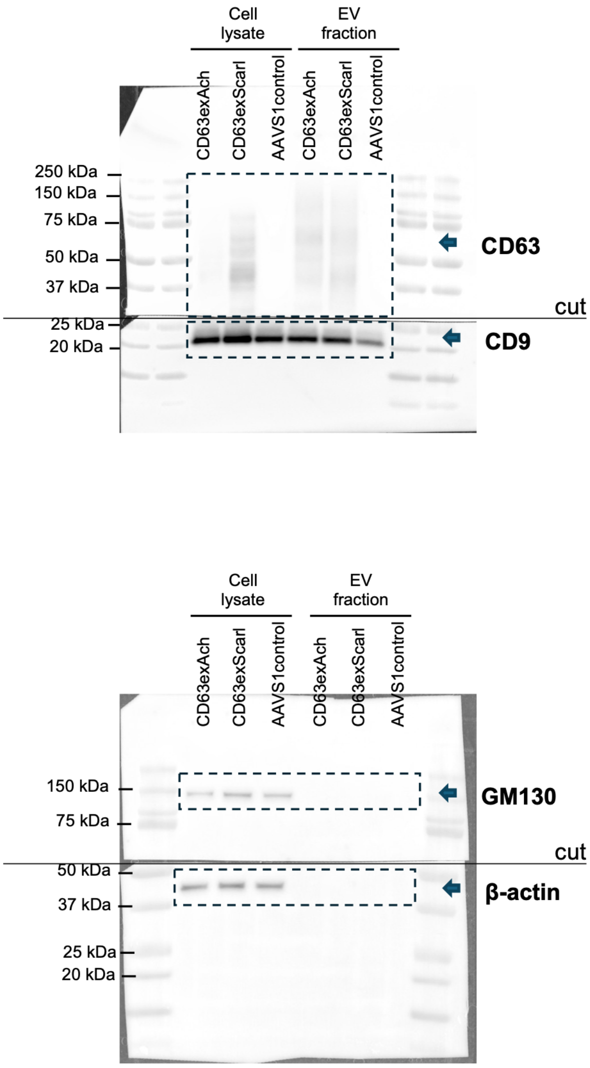


**Fig. S3 Uncropped full-length pictures of western blotting membranes in Fig. 1.**

Membranes were often cut to enable blotting for multiple antibodies. The areas outlined by black dotted boxes in the raw blots correspond to the cropped images shown in Fig. 1D.

Fig. S4

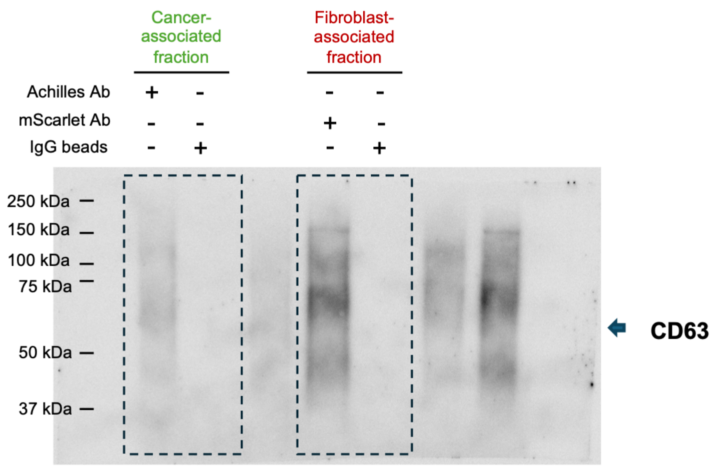


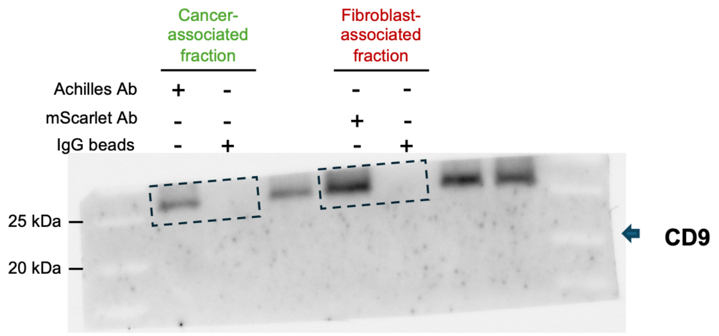


**Fig. S4 Uncropped full-length pictures of western blotting membranes in Fig. 4.**

The areas outlined by black dotted boxes in the raw blots correspond to the cropped images shown in Fig. 4B.

**Video S1. Dynamic movement of EV-Tracer-positive puncta in direct cancer–fibroblast co-culture.**

44As3 CD63exAch cancer cells and iNF-58 CD63exScarl fibroblasts were directly co-cultured and imaged every 5 min for 145 min (30 time points) using a BZ-X810 fluorescence microscope. Achilles and mScarlet signals are displayed in yellow and red, respectively. The sequence corresponds to Fig. 3E and shows dynamic changes in the position of reporter-positive puncta within and between neighboring cellular regions. The imaging data do not independently establish definitive EV transfer or cell of origin.
